# Predicting the Evolutionary Consequences of Trophy Hunting on a Quantitative Trait

**DOI:** 10.1101/127555

**Authors:** Tim Coulson, Susanne Schindler, Lochran Traill, Bruce E. Kendall

## Abstract

Some ecologists suggest that trophy hunting (e.g. harvesting males with a desirable trait above a certain size) can lead to rapid phenotypic change, which has led to an ongoing discussion about evolutionary consequences of trophy hunting. Claims of rapid evolution come from the statistical analyses of data, with no examination of whether these results are theoretically plausible. We constructed simple quantitative genetic models to explore how a range of hunting scenarios affects the evolution of a trophy such as horn length. We show that trophy hunting does lead to trophy evolution defined as change in the mean breeding value of the trait. However, the fastest rates of phenotypic change attributable to trophy hunting via evolution that are theoretically possible under standard assumptions of quantitative genetics are 1 to 2 orders of magnitude slower than the fastest rates reported from statistical analyses. Our work suggests a re-evaluation of the likely evolutionary consequences of trophy hunting would be appropriate when setting policy. Our work does not consider the ethical or ecological consequences of trophy hunting.

---

Trophy hunting that is well managed, and based on robust monitoring protocols, can be a useful conservation tool in areas where there is increasing demand for land from growing human populations (Di Minin et al., 2016; Lindsey et al., 2006). The logic of the approach is that selectively hunting a small proportion of males with large horns, antlers, or body size, will have few ecological and evolutionary consequences because species with sexually selected characters usually exhibit a polygynous mating system in which males are not limiting (Dickson et al., 2009; Milner-Gulland and Mace, 1998). However, a debate on the ethics, use, and consequences, of trophy hunting is underway (Lindsey et al., 2016; Nelson et al., 2016; Ripple et al., 2016), including an ongoing fast or slow evolution discussion on hunted bighorn sheep (*Ovis canadensis*; Coltman et al., 2003; Pigeon et al., 2016; Traill et al., 2014). We contribute further to the trophy hunting debate by constructing and analysing general quantitative genetic models of the effect of trophy hunting on phenotypic evolution.

Proponents of trophy hunting argue that selling the rights to selectively hunt individuals with desirable attributes is a useful way to raise money (Rodríguez-Muñoz et al., 2015). The argument is that if wildlife populations can be moneterized, they have value, and this worth makes the area in which the population lives more easily protected from competing land use interests (Lindsey et al., 2007). Profit generated from hunting can be invested in conservation, habitat improvement or in local communities, and any ecological and evolutionary consequences of selective hunting on males is likely to be a small cost worth paying (e.g., Crosmary et al., 2015).

Those opposed to the approach argue either that trophy hunting is unethical, or that money raised from trophy hunting rarely gets invested in local communities or in conservation. For example, in Africa, monies raised from selling hunting rights can get subsumed into government coffers, and profits made by outfitters do not always make it back to the local area or communities (Lindsey et al., 2014). In addition, the ecological outcomes of hunting may be negative: in East Africa, unregulated trophy hunting influenced a localized extirpation of lion (*Panthera leo*) populations (Packer et al., 2011), and unethical lion hunting practices in Hwange National Park in Zimbabwe resulted in 72% of research animals being killed, including 30% of males < 4 years old that had yet to breed (Loveridge et al., 2007). Furthermore, hunting may lead to evolution of selected traits as has frequently been speculated for some sheep populations (Douhard et al., 2016; Festa-Bianchet et al., 2014; Pigeon et al., 2016).

One reason why the ecological and evolutionary consequences of trophy hunting have received recent interest is that biologists have found that evolution can be observed on ecological timescales (Hairston et al., 2005). This has spawned the field of eco-evolution (Schoener, 2011). There is compelling empirical evidence of rapid, joint phenotypic and ecological change from a number of systems (e.g., Hairston et al., 2005; Ozgul et al., 2010), but evidence of genetic change is much less widespread (Yoshida et al., 2003), partly because it is harder to demonstrate. Quite frequently, phenotypic change is attributable to evolution without supporting evidence of genetic change (Hendry, 2016), or without examining whether the rates of evolutionary change reported are theoretically plausible (Coltman et al., 2003).

Coltman et al. (2003) report rapid phenotypic change in the face of hunting that was attributed to evolution. Based on longitudinal data for the Ram Mountain bighorn population in Canada Coltman et al. (2003) used statistical quantitative genetics to argue that selective hunting of, on average, 2 rams/year from a population of, on average, approximately 70 bighorn sheep resulted in a 30% decline in horn size over 5 generations. In a second paper Pigeon et al. (2016) reported a new analysis that supports these claims. These papers (Coltman et al., 2003; Pigeon et al., 2016) have become influential as opponents of trophy hunting argue that the activity has rapid detrimental consequences on hunted populations. However, no papers have yet examined whether the rates of change observed by Coltman et al. (2003) and Pigeon et al. (2016) are plausible using the quantitative genetic theory that motivated their statistical analyses, even though skepticism has been raised as to whether the phenotypic changes observed can be attributed to evolution (e.g., Traill et al., 2014).

We developed novel, general theory to examine the likely evolutionary consequences of selective harvesting on a single sex in a sexually reproducing species. We worked in the quantitative genetics framework because the genetic architecture of trophy traits is rarely known (e.g., Kruuk et al., 2002). We start with a brief summary of quantitative genetic theory that motivated our models, and which is widely used to examine the evolution of phenotypic traits of unknown genetic architecture in free-living populations (Merilä et al., 2001). We then describe the models we used, along with the parameter values we selected.

## METHODS

We use the following notation. Expectations and variances of the distribution of *N*(*x*,*t*) are denoted 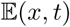 and *V*(*x, t*) respectively. A subscript, either of *f* or *m*, is used to identify distributions or moments of distributions taken over only females or males respectively. If this subscript is absent, the distribution is taken over both sexes. We use a superscript *R* to identify distributions, or moments of distributions, that have been operated on by selection.

### A Quantitative Genetic Primer

Quantitative genetics assumes that an individual’s phenotype *Ƶ* consists of the sum of various components. These components include a breeding value 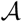 and the environmental component of the phenotype ***Ɛ***, with contributions from epistasis and non-additive genetic effects also sometimes included in the sum (Lynch and Walsh, 1998). Only 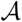 and ***Ɛ*** are considered here. An individual’s breeding value describes the additive genetic contribution to its phenotypic trait value. But what does this mean?

If alleles at a locus have an additive effect on a phenotypic trait, each allele can be assigned a value that describes the contribution of that allele (in any genotype at that locus) to the phenotype. For example, consider a bi-allelic locus with 3 genotypes, *aa, aA* and *AA*. Allele *a* has a value of 1g and allele *A* a value of 2g. The breeding value of each genotype to body mass will be: *aa* =1 + 1 = 2, *aA* = 1 + 2 = 3, and *AA* = 2+2 = 4. Breeding values can be summed across genotypes at different loci to generate breeding values for multi-locus genotypes. Under the additivity assumption, the dynamics of breeding values is identical to the dynamics of alleles; this is the not always the case when non-additive genetic processes like heterozygote advantage and epistasis are operating (Falconer, 1975).

Many applications of quantitative genetics use the infinitesimal model (Fisher, 1930). This assumes that an individual’s breeding value for a phenotypic trait is made up from independent contributions from a large (technically infinite) number of additive genotypes, each making a very small contribution to the phenotypic trait. There is no interaction between alleles at a locus (dominance) or interactions between genotypes at different loci (epistasis).

In additive genetic models used to predict evolutionary change, it is usually assumed that ***Ɛ*** is determined by developmental noise. An individual’s environmental component can be considered as a random value drawn from a Gaussian distribution with a mean and a constant variance: norm(0, *V*(***Ɛ**, t*)). 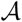 and ***Ɛ*** are consequently independent. Thus,

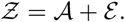

The distribution of breeding values is also assumed to be Gaussian, and

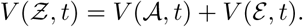

These assumptions mean that, on average, the breeding value can be inferred from the phenotype – the phenotypic gambit. The aim of statistical quantitative genetics is to correct the phenotype for nuisance variables so the phenotypic gambit assumption is appropriate for the corrected phenotype (Lynch and Walsh, 1998).

Next, quantitative genetic theory makes the assumption that the mean of 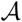 among parents is equal to that in offspring: e.g., 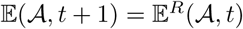. In 2-sex models this requires that the expected value of 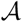 in an offspring is the mid-point of the breeding value of its parents. Given this assumption,

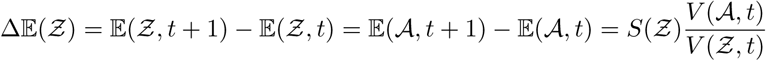
where 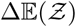 is the difference in the mean of the phenotype between the offspring and parental generations, *S*(*Ƶ*) is the selection differential on *Ƶ* and 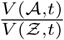 the heritability (*h*^2^) of a trait, and *t* represents generation number. The selection differential describes the difference in the mean value of the character between those individuals selected to reproduce and the entire population prior to selection (Price, 1970). Equation () is the univariate breeders equation (Falconer, 1975).

If all assumptions of the univariate breeders equation are met, it will accurately predict evolution of a trait assuming that the selection differential and the additive genetic and phenotypic variances have been appropriately estimated. One exception where it can fail is if there are genetically correlated characters that have not been measured, and which are under selection (Lande and Arnold, 1983).

Lande and Arnold (1983) developed a multivariate form of the breeders equation that states:

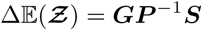
where 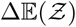 is a vector describing change in the mean of each of the phenotypic traits from the parental to the offspring generation, ***S*** is a vector of selection differentials on each character, ***G*** is a genetic variance-covariance matrix, and ***P*** is a phenotypic variance-covariance matrix. If 2 traits are genetically correlated, and both are under selection, to understand how 1 of the traits evolves it is necessary to understand how the 2 traits are genetically and phenotypically correlated, and how strong selection is on each of the traits.

In both the univariate and multivariate breeders equations, the selection differentials capture total selection (Lande and Arnold, 1983). This means that both equations accurately capture selection on the trait(s) even in the presence of unmeasured genetically correlated characters. Genetically correlated characters influence predictions of evolution in the breeders equations through their impact on estimates of the heritability (in the univariate case) and the ***G*** matrix (in the multivariate case).

A limitation of the breeders equation is it is not dynamically sufficient – it should not be used to make predictions across multiple generations, particularly when evolution is sufficiently strong that it alters genetic variances and covariances (Lande and Arnold, 1983). To construct a dynamic model, it is either necessary to make assumptions about the genetic variance (it is sometimes assumed to be constant: (Lande, 1982)) or to track the dynamics of the entire distributions of 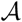 and ***Ɛ*** (Coulson et al., 2017) or 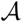 and *Ƶ* (Barfield et al., 2011; Childs et al., 2016). We model the dynamics of 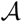 and ***Ɛ***.

### A Generic Model to Explore the Effects of Trophy Hunting on Evolution

We developed a 2-sex, dynamic, quantitative genetic model to explore how hunting on one sex influences phenotypic evolution. We iterate the population forwards on a per-generation time step.

We assume that in the absence of hunting, the trophy is not under selection in either sex and is consequently not evolving. This provides us with a baseline scenario in the absence of hunting with which to compare results from a range of hunting scenarios. We define a bivariate distribution *N*(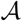, ***Ɛ**, t*) of breeding values 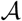 and the environmental component of the phenotype ***Ɛ*** in generation *t*. At time *t* = 0 we assume the distribution *N*(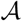, ***Ɛ**, t*) is bivariate normal with means ***μ*** and (co)variances **Σ**. The 2 components of ***μ*** are 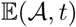 and 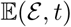 at *t* = 0. **Σ** is a variance-covariance matrix,

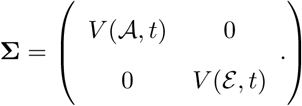

Variances can be chosen to determine the heritability *h*^2^ at time *t* = 0.

We assume that males and females have the same distribution of phenotypes and breeding values at birth, and that the birth sex ratio is unity: *N_f_* (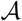, ***Ɛ**, t*) = *N_m_*(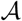, ***Ɛ**, t*) = 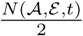.

Next we impose selection. There is no direct selection on females and the number of recruits they produced is set to 2, the replacement rate, to ensure the female population remains the same size over time and the population growth rate λ = 1. This assumes males are not limiting. The distribution of females selected to reproduce is consequently 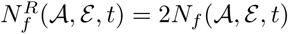. The same function for males is used in the absence of hunting.

When males are selectively hunted, we remove individuals from the distribution before assigning male reproductive success. We then scale the resulting distribution of males to be the same size as the distribution of females. For example, if all males of above mean trophy size are culled, the matings they would have had are redistributed across those males that were below the mean trophy size and not hunted. In the case of a Gaussian distribution of the trophy, their lifetime reproductive success would increase proportionally to the number of males culled. The proportion *p* is calculated and the post-selection distribution 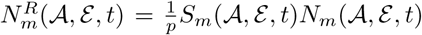 is calculated where *S_m_*(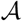, ***Ɛ**, t*) is the function describing selection on the male trophy. The distribution 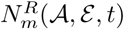 is the distribution of the components of the phenotype of those males selected to be fathers.

We impose selection on males by culling a proportion α of individuals that are above average size,

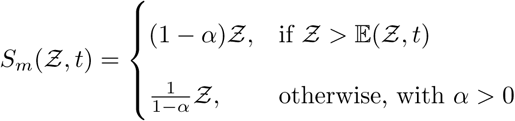

This generates a distribution of fathers 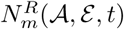 that is equal in size to the distribution of mothers 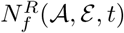.

We now have distributions of maternal and parental characters that are the same sizes and sufficient for the female population to replace itself with some males reproducing with multiple mothers. We assume random mating and calculate the distribution of parental midpoint breeding values *N^R^*(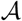,*t*) by convolving 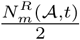 with 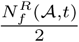. To generate the distribution of offspring breeding values, we convolve this distribution with a distribution of the segregation variance, defined as a Gaussian distribution with a mean of 0 and a variance equal to half the additive genetic variance of the distribution 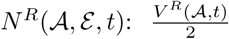 (Barfield et al., 2011). Effects of increases in the additive genetic variance via mutation, or from other sources of genetic variation being converted to additive genetic variance, can be captured by increasing the size of the segregation variance. Finally, we generate a distribution of the environmental component of the phenotype for each value of 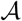 in the offspring distribution that is proportional to a Gaussian distribution with a mean of 0 and an environmental variance that is the same as that in the previous generation. We now have the bivariate distribution of the components of the phenotype in offspring *N*(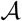, ***Ɛ**, t* + 1).

Taken together this gives the following recursion,

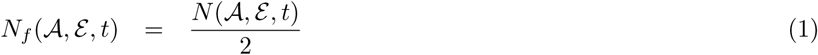

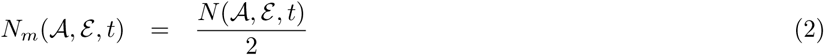

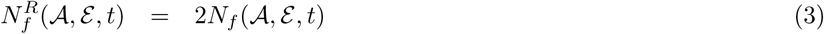

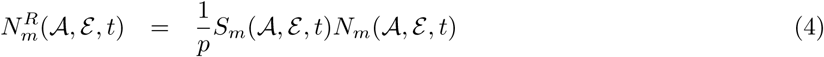

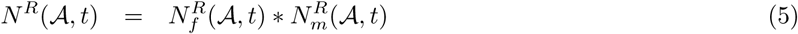

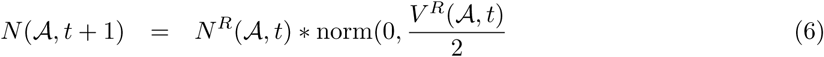

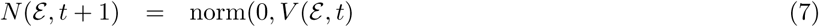

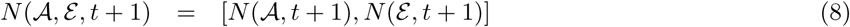

### Analysis of the Multivariate Breeders Equation

When evolutionary predictions fail to match observation, the existence of correlated unmeasured characters is often assumed (Merilä et al., 2001). However, the potential impact of correlated characters on evolution assuming selection differentials have been appropriately measured is rarely investigated. We used the multivariate breeders equation to examine how such characters can influence evolution, and in particular, whether they can generate rapid evolution in directions opposite to those predicted by selection differentials which measure total selection on a trait (Lande and Arnold, 1983).

We assume 2 traits *Ƶ*_1_ and *Ƶ*_2_. We predict 1 generation ahead, so we do not use *t* for time to simplify notation. We define bivariate Gaussian distributions of the traits’ breeding values 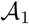 and 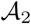 (norm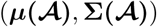) and environmental components of the phenotype (norm(***μ*(***Ɛ*****), **Σ**(*****Ɛ*****))). From this we construct a bivariate Gaussian distribution of the phenotype norm(***μ***(***Ƶ***), **Σ**(***Ƶ***)) = norm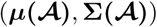 + norm(***μ*(***Ɛ***), Σ(***Ɛ***)**).

We now impose selection on the phenotype with the following fitness function *W*(*Ƶ, t*) = *β*_0_ + *β*_1_*Ƶ*_1_+*β*_2_*Ƶ*_2_. We estimate selection differentials on the 2 phenotypic traits as ***S*** = **Σ**^−1^(***Ƶ***)***β*** where ***β*** = (*β*_1_,*β*_2_)*^T^* where *T* is the vector transpose and ***S*** is a vector containing the selection differentials *s*_1_ and *s*_2_. We also calculate the univariate fitness functions 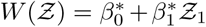 and 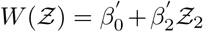 using methods from instrumental variable analyses (Coulson et al., 2017; Kendall, 2015). From these functions, we calculated the univariate selection differential 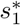 and 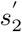. We calculate univariate heritabilities using the relevant additive genetic variances and phenotypic variances for each trait. We then compare predictions of evolutionary change between the multivariate breeder’s equation and the two univariate breeder’s equations.

### Model Parameters

We set ***μ*** = [70 cm, 0]. The value of 70 cm is approximately the mean horn length of 4-year old rams reported by (Coltman et al., 2003, Figure 2) at the start of their study. The value of zero is the mean of the environmental component of the phenotype as is usually assumed in quantitative genetics (Falconer, 1975).

To explore the effects of hunting on the evolution of a trophy, we ran simulations with a range of initial genetic, environmental and phenotypic variances. We conducted simulations to demonstrate the effects of altering the additive genetic variance and the total phenotypic variance. For example, the simulations reported in fig. 1 and fig 2. both have identical initial additive genetic variances of *V*(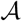, 1) = 3 but they have different environmental variances of *VƐ*, 1 = 2 and *VƐ*, 1 = 0.1 respectively. The simulations reported in Fig. 3 demonstrate the effect of increasing the phenotype variance by increasing the additive genetic variance compared to those simulations reported in Fig. 1 and Fig. 2: *V*(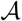, 1) = 5 and *V*(***Ɛ***, 1) = 2.

**Figure 1.**
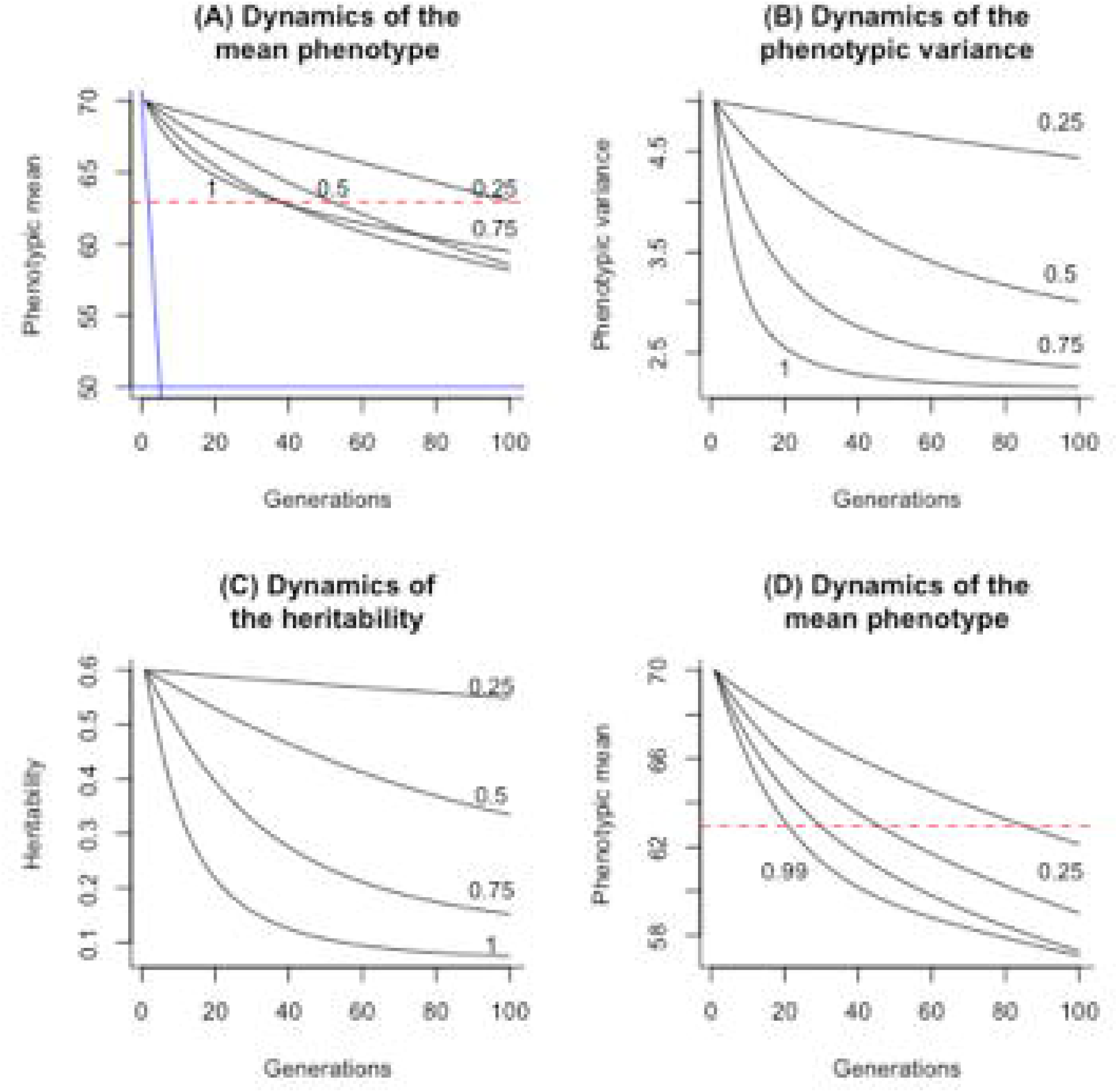
The effect of different trophy hunting regimes on the dynamics of the phenotype and the heritability. The dynamics of the mean (A), the variance (B) and the heritability (C) all depend upon the proportion of males of above average trophy (e.g., horn) size that are culled (numbers next to lines). In (A) the red horizontal line represents 1.96 standard deviations from the initial mean trophy size. We selected the starting mean phenotypic value in (A) to be the same as that reported by Coltman et al. (2003). The blue horizontal line is the mean phenotype Coltman et al. (2003) reported 5 generations later. The near vertical blue line represents the rate of the change in the phenotypic mean they report. The line can be compared with the lines from our simulations. In these simulations, the initial additive genetic variance was set at 3.0, and the environmental variance at 2.0. We also report the dynamics of the mean phenotype when 25% of above average trophy sizes are harvested as a function of increasing additive genetic variance and the heritability (D). In each of the four simulations reported in (D) we set the initial phenotypic variance at 5 by using values for the initial additive genetic variances as (4.99,3.75,2.5,1.25) and for the environmental variances as (0.01,1.25,2.5,1.75). These give initial heritabilities of 0.99,.075,0.5 and 0.25 (values next to the lines).

**Figure 2.**
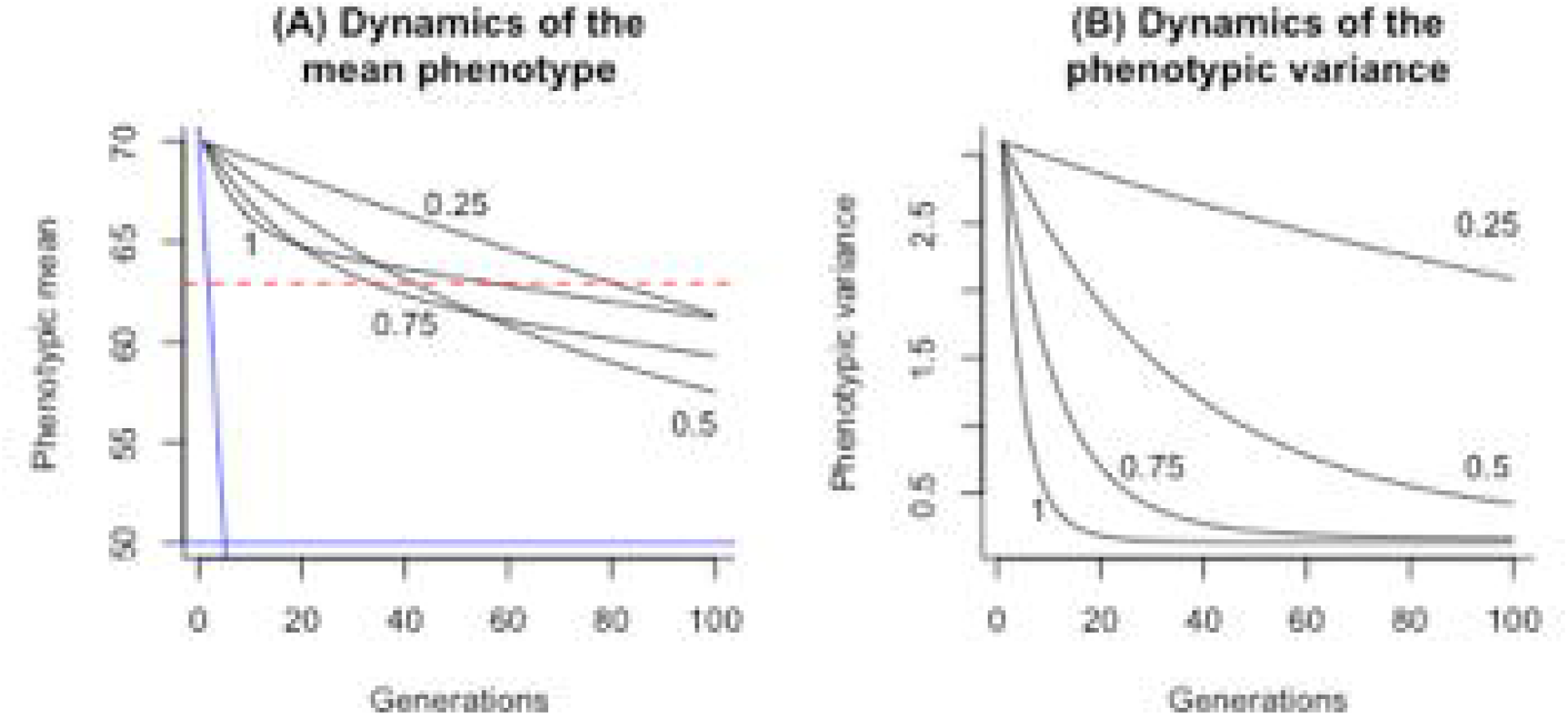
The effect of different trophy hunting regimes on the dynamics of the phenotype. The dynamics of the mean (A), and the variance (B) for cases when the phenotype is determined almost entirely by the additive genetic variance. In each simulation the initial additive genetic variance was set to 3.0 and the environmental variance to 0.1. The blue horizontal line is the mean phenotype Coltman et al. (2003) reported 5 generations later. The near vertical blue line represents the rate of the change in the phenotypic mean they report. The line can be compared with the lines from our simulations.

We also examined the consequences of injecting additional genetic variance into the population at each time step by setting the segregation variance to the constant initial value chosen at the beginning of the simulation. For all parameter sets, we explored the effect of removing 25%, 50%, 75%, and 100% of males of above average horn size (e.g., *α* = [0.25,0.5,0.75,1]).

To demonstrate how correlated characters affect phenotypic evolution over a single generation, we ran a number of simulations of the multivariate breeder’s equation. In each simulation we set *ω* = 0.3 + 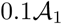 + 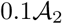 and ***μ*(*Ƶ*)** = (6, 6). These values are arbitrary in that any values could be used to reveal the effects we demonstrate. We then ran 12 simulations. In each simulation *V*(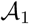, *t* = 0) = 2 and *V*(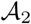, *t* = 0) = 2. We then examine 3 genetic covariance structures within 4 different distributions of the environmental components of the phenotype. The first assumes no genetic covariance, the second a negative genetic covariance of-1.41 and the third a positive genetic covariance of 1.41. We chose the second and third values because they are the 2 limits that the covariance can take to ensure the variance-covariance matrix is positive-definite. The 4 distributions of the environmental components of the phenotype are selected such that phenotypic variances and covariances are dominated by the additive genetic variances and covariances, and for cases where approximately half of the phenotypic variances and covariances are attributable to the additive genetic variances and covariances. We then explored the effects of positive and negative covariances between the environmental components of the phenotypes on evolutionary dynamics.

## RESULTS

Selective trophy hunting led to an evolutionary response in all of our simulations (Fig. 1-3). In our initial simulation with a starting heritability of 0.6, the phenotypic mean declined from a initial value of 70 to between 57 and 62.5 depending upon the proportion of the population culled. There was relatively little difference in the mean phenotype after 100 generations when 50%, 75%, or 100% of males of above average trophy value were harvested; all simulations achieved a decline from 70 to 57 over 100 generations. In contrast, evolution was notably slower when only 25% of above average trophy sizes were culled per generation (Fig. 1(A). The phenotypic variation and heritability showed similar rates of change. This is expected because variation in the environmental component of the phenotype at birth is constant across generations. The rate of loss of phenotypic variation and decline in the heritability scaled with harvesting rate (Fig. 1(B,C)). When all males above the mean trophy value were harvested, additive genetic variance was initially rapidly eroded, before starting to decline more slowly. This change was reflected in the dynamics of the phenotypic variance (Fig. 1(B)). These rates of change in the variance affected the dynamics of the mean phenotype. Although the initial rate of evolution correlated with harvesting pressure, over the course of 100 generations evolution was fastest when 75% of above average males were harvested. None of our scenarios predicted phenotypic change at the rate reported by Coltman et al. (2003). In our initial simulations it took between 40 and 100 generations before the mean phenotype evolved to a value that would be significantly different from its initial value (regardless of sample size). Finally, altering the initial heritability by reducing the initial additive genetic variance slowed the rate of evolutionary changed as expected. In contrast, as the additive genetic variance and consequently heritability increased, so too did the rate of evolution (Fig. 1(D)).

**Figure 3.**
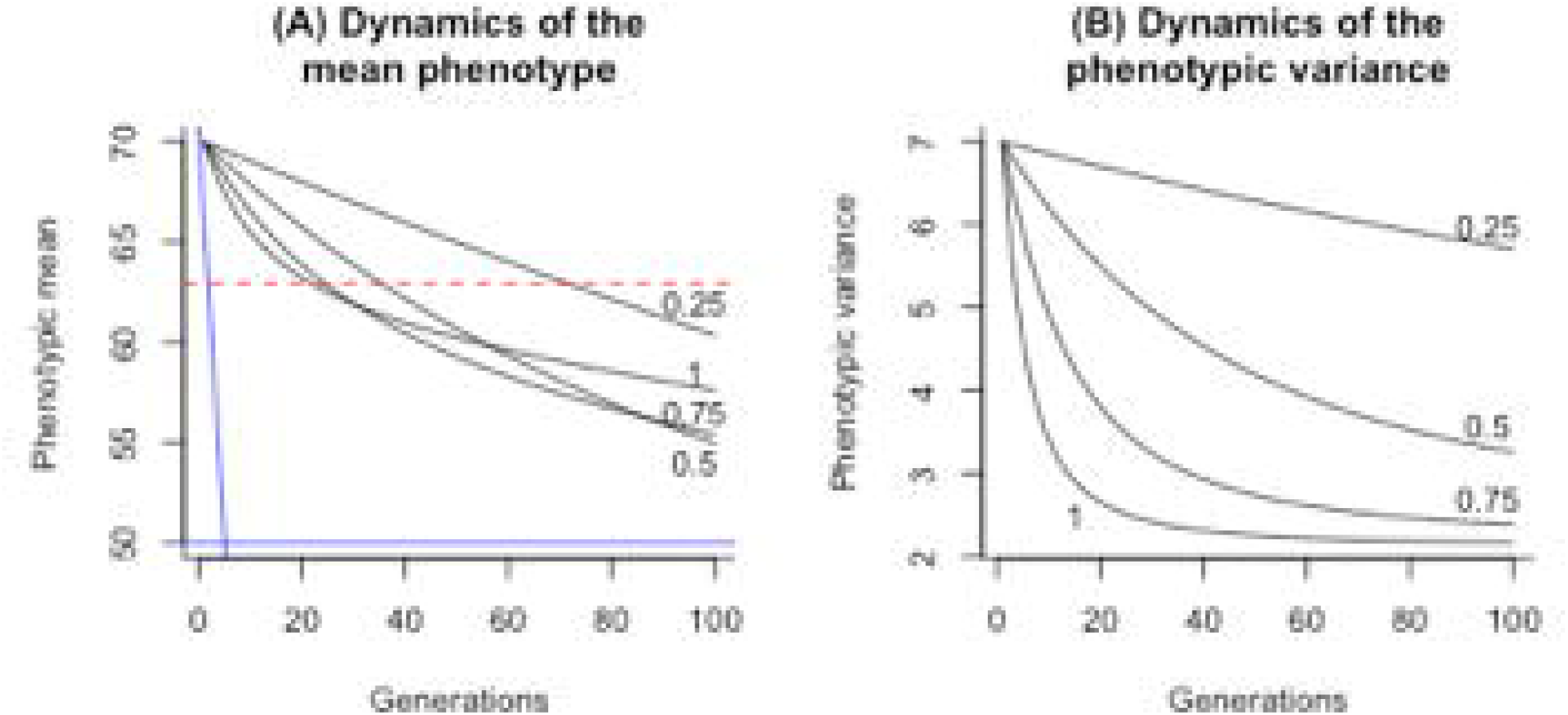
The effect of different trophy hunting regimes on the dynamics of the phenotype. The dynamics of the mean (A) and the variance (B) to demonstrate the effect of a high heritability and large phenotypic variance. In each simulation the initial additive genetic variance was set to 5.0 and the environmental variance to 2.0. The blue horizontal line is the mean phenotype Coltman et al. (2003) reported 5 generations later. The near vertical blue line represents the rate of the change in the phenotypic mean they report. The line can be compared with the lines from our simulations.

In our second simulation, we increased the initial heritability by reducing the environmental variation. This had a relatively small impact on the rates of evolution (Fig. 2(A)), although the reduction in the phenotypic variance (Fig. 2(B) did reduce rates of evolution at the highest levels of off-take (Fig. 2(A). Increasing the additive genetic variance, and consequently the phenotypic variance, also increased rates of evolutionary change slightly (Fig. 3(A,B)), although rates of evolution were still between 1 and 2 orders of magnitude slower than reported by Coltman et al. (2003) and Pigeon et al. (2016). The time series of selection differentials estimated across males and females for these simulations are given in Fig. S1.

In all simulations, setting the segregation variance to a constant value generated linear selection because selection does not rapidly erode the additive genetic variance (Fig. S2). However, even when all males of above average horn size are culled, the rate of evolution is still > 5 times slower than that reported by Coltman et al. (2003).

We next compared evolutionary dynamics predicted by the univariate and bivariate breeders equation to examine whether correlated characters could lead to rapid evolution in the opposite direction to selection, or to evolutionary stasis. The degree of correlation between 2 characters increased the rate of evolution when the sign of the phenotypic covariance (−/+) was the same as the sign of the product of the selection differentials on each trait (Fig. 4(A)-(D)). As the proportion of phenotypic variation attributable to additive genetic variation tended to unity, predictions from the univariate and bivariate breeders equation converged (Fig. 4(A)). Similarly, although not reported, at the other limit, as the proportion of phenotypic variance attributable to additive genetic variance tended to zero, no evolution was predicted by either the univariate or bivariate breeders equation and predictions converged. Departures between the 2 equations were greatest when intermediate proportions of the phenotypic variances and covariances were attributable to the additive genetic variances and covariances (Fig. 4(B)-(D)). Both additive genetic covariances, and covariances in the environmental component of the phenotype, could lead to divergence between the univariate and bivariate breeders equation (Fig. 4(B)-(D)).

**Figure 4.**
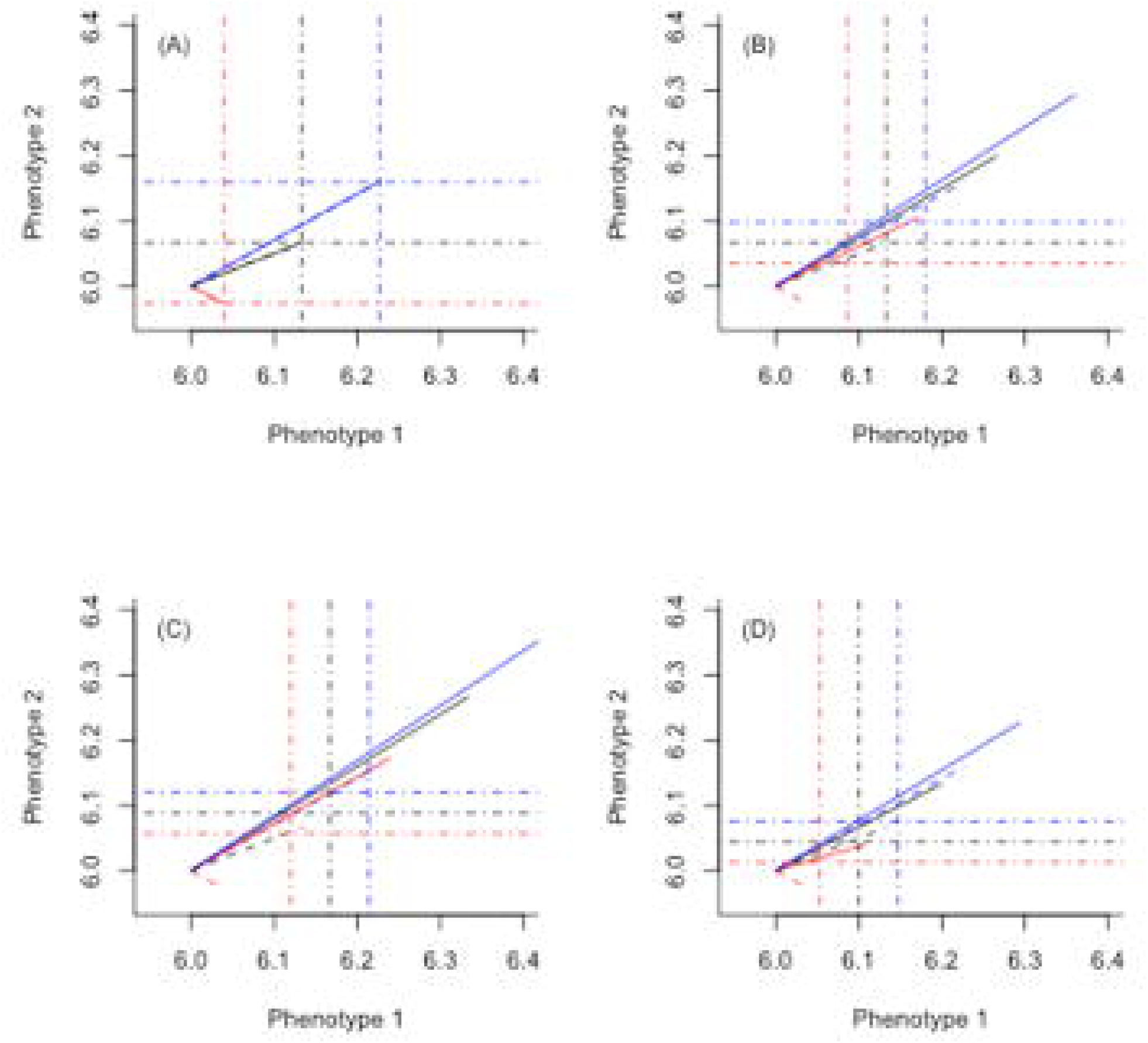
A comparison of the dynamics of the multivariate and univariate breeders equation for different degrees of additive genetic and environmental variances and covariances. Each figure reports 3 simulations: no genetic covariance (black lines), strong positive genetic covariances that reinforce selection (blue lines), and strong negative genetic covariances that oppose selection (red lines). Solid lines represent selection differentials on each trait and dotted lines represent responses to selection. Horizontal and vertical dot-dashed lines show predictions of evolution from the univariate breeders equation for each trait. The farther the right hand end of the dashed lines are from the intersection of the horizontal and vertical dot-dashed lines, the greater the disparity between predictions from the univariate and multivariate breeders equation. We simulated that all phenotypic variation is attributable to genetic (co)variances (A), approximately half of phenotypic variance is attributable to additive genetic variance (B), and the effect of a positive (C) and negative (D) covariance in the environmental components of the phenotypes on rates of evolution. The genetic and environmental (co)variance used in each simulation can be found in Table S1.

Although covariances in **Σ(***Ɛ***)** and **Σ**(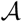) could affect rates of evolution, when selection differentials were large, covariances could not generate stasis or lead to evolution in the opposite direction to that predicted by selection (Fig. 4, blue lines). However, as selection got weaker, correlated characters could prevent selection, and even lead to very small evolutionary change in the opposite direction to that predicted by evolution (Fig. 4, red lines). However, effect sizes were small and would be challenging to detect without large quantities of data.

## DISCUSSION

Our simulations show that selective harvesting can alter the evolutionary fate of populations, and can result in declines in trophy size. However, even under intensive trophy hunting, it is expected to take many tens of generations before the mean trophy size has evolved to be significantly smaller than it was prior to the onset of selective harvesting (see also Mysterud and Bischof, 2010; Thelen, 1991). Our results also show that although correlated characters can have impacts on phenotypic evolution, they cannot be invoked to explain rapid phenotypic change in the opposite direction to that predicted from univariate selection differentials.

Our models are kept deliberately simple and make a number of assumptions. First, we iterate the population forwards on a per-generation step. This means there is no age structure, and that a single breeding value determines trophy size throughout life. For some traits there is evidence of age-specific breeding values (Wilson et al., 2005), and these could influence evolutionary rates (Lande, 1982). Males are typically shot once they have reached adulthood, which means direct selection via hunting does not occur in younger ages. The indirect effect of trophy hunting at older ages on phenotypes and fitness at younger ages is determined by genetic correlations across ages. As we show in our analysis of the multivariate breeders equation, evolution is most rapid when the genetic correlations are close to the limit and align with the direction of selection. Given trophy sizes typically experience positive selection at all ages (Coltman et al., 2002; Preston et al., 2003), this means that the rate of evolution will be greatest when genetic correlations are close to unity. At the limit, this would mean that the same breeding value would determine trophy size throughout life – an assumption of our model. Our model consequently likely predicts faster rates of evolution than would be predicted from a model with age-structured breeding values and the same selection regime that we assume.

A second assumption we make is that the trait is not subject to selection before selective harvesting is imposed. Trophy size positively correlates with fitness in species that are not harvested (Preston et al., 2003). Trophies may consequently be expected to be slowly evolving to be larger in the absence of selective hunting. If that were the case, then the effect of trophy hunting would have to be greater than in our models to lead to evolution of smaller trophies at the rates we report. This is because selective harvesting would have to counteract evolution for larger trophies in the absence of harvesting, before then leading to a reduction in trophy size. Our model would over-estimate the evolutionary impact of trophy hunting in such a case.

Males in sexually dimorphic species with trophies form dominance hierarchies (Pelletier and Festa-Bianchet, 2006). If a dominant male with large trophies is shot, it may be reasonable to assume that surviving males with large trophies that are towards the top of the dominance hierarchy would secure the re-productive success the shot male would have enjoyed. We do not model this process. Instead, we redistribute the reproductive success across all remaining males. This egalitarian redistribution of reproductive success likely exaggerates the evolutionary consequences of trophy hunting because individuals with small trophies are benefiting from those with large trophies being shot. Our model, although simple, has consequently been formulated to likely exaggerate the consequences of trophy hunting on trophy evolution.

When predictions from simple models like ours fail to match with observation, the existence of genetically correlated unmeasured characters is often invoked as an explanation (Merilä et al., 2001). Changing the degree of generic covariation between two characters can significantly alter selection differentials on both characters (e.g., Fig. 4). However, this does not mean that the failure to measure a correlated character will lead to incorrect estimates of a selection differential on a trait. In fact, the failure to measure a correlated character will have no impact on the estimate of a selection differential of a focal character (Coulson et al., 2017; Kingsolver et al., 2001; Lynch and Walsh, 1998). Estimates of selection differentials on a univariate character will consequently always give an upper limit on the rate of evolution of a character that conforms to the assumptions of the phenotypic gambit.

Genetic and environmental covariation with unmeasured characters can affect the response to selection. The effect is most likely to be strongest when characters have heritabilities in the vicinity of 0.5 and co-variances are close to their limits. The further from this proportion that variance and covariances get, the less biased predictions of evolution in the presence of unmeasured correlated characters becomes. Large covariances that act to reduce the strength of selection can lead to low rates of evolutionary change in the opposite direction to selection, but the effect is small and could only be detectable in very large data sets. We consequently conclude that if the phenotypic gambit is assumed and significant selection on a trait is observed, then unmeasured correlated characters can act to slow, or increase, rates of evolution compared to those predicted by the univariate breeders equation, but they cannot result in evolutionary change that is greater than the univariate selection differentials, or lead to evolutionary stasis. We conclude that although our models on the effect of hunting on a trophy are simple, they will not be too wide of the mark, particularly for large initial heritability for a trophy.

Although our models are simple, they provide some novel insights. In particular, our strongest selection regimes result in initial increased rates of evolution. However, they erode the additive genetic covariance more quickly than less stringent hunting regimes, rapidly slowing the rate of evolution. Over longer periods, evolutionary rates are highest at intermediate rates of hunting compared to higher hunting rates. These results show how important it is to track the dynamics of the additive genetic variance when predicting evolution in the face of strong selection over multiple generations (see also Barfield et al., 2011; Childs et al., 2016; Coulson et al., 2017; Lande, 1982). Assuming a constant additive genetic variance in the face of strong selection would lead to predictions of elevated rates of evolution over multiple generations.

In most of our simulations we assume that the directional selection we impose erodes the additive genetic variance as is often assumed in quantitative genetics (Falconer, 1975). We do this by constraining the segregation variation to be equal to half the additive genetic variance among parents (Barfield et al., 2011; Childs et al., 2016). However, we also relax this assumption by maintaining a constant segregation variance that is not eroded in the face of selection. This mimics processes, including mutation, that generate additive genetic variance. By doing this we linearize the longer-term response to selection, such that evolution continues to alter the trait value at a greater evolutionary rate over a longer period of time than is possible when selection erodes the additive genetic variance. However, even under these circumstances, statistically significant evolution is predicted to take between 10 and 20 generations even under strong selection when all males of above average horn size are culled.

What do our results contribute to bighorn sheep management? Their primary contribution is to suggest that very fast phenotypic change of quantitative characters that is sometimes observed in these populations cannot be due to rapid evolution, at least not under the assumptions of quantitative genetics, for 2 reasons. First, the upper rates of change reported (Coltman et al., 2003) are approximately 2 orders of magnitude faster than models of intensive selective harvesting can achieve. Second, the traits that are hypothesised to evolve, horn length and body size, are subject to positive selection at some ages, even in the presence of harvesting (Traill et al., 2014), yet body and horn size have become smaller (Coltman et al., 2003). Unmeasured correlated characters cannot explain this. So what causes the rapid phenotypic change that is sometimes observed? There are a number of possibilities.

First, the environment may have deteriorated rapidly, leading to a change in the mean of the environmental component of the phenotype (Kruuk et al., 2002; Merilä et al., 2001), perhaps in a similar manner as reported in a desert bighorn sheep population (*Ovis canadensis nelsoni*; Hedrick, 2011). Second, the phenotypic gambit on which statistical quantitative genetic analyses are based, may be violated (Hadfield et al., 2007). This could occur if genotype-by-environment interactions, dominance variation or epistasis have contributed to the observed phenotypic trends (Falconer, 1975). Quantitative genetics theory and empirical methods exists to deal with each of these processes (Lynch and Walsh, 1998), but statistical methods to estimate these processes either require large population sizes or additional data that may not be available for this population. Third, the association between body size and horn length and fitness may not be causal (Merilä et al., 2001), but both may reflect an individuals ability to extract resources from the environment. Individuals that are good at doing this grow to large sizes, produce large trophies, and have high fitness. If the ability to extract resources from the environment is not determined by a simple additive genotype-phenotype map, then neither will be the association between body size and horn length and fitness.

Although our models reveal that very rapid evolution attributable to selective hunting is not a plausible explanation for the observed phenotypic declines, our models are not parameterized for bighorn sheep. Ideally the theoretical quantitative genetic approach we use here and in Coulson et al. (2017) should be parameterized for bighorn sheep before any management recommendations are made. The only data set we are aware of that may be sufficient to parameterize models within our framework are from the bighorn sheep population at Ram Mountain (Coltman et al., 2003; Pigeon et al., 2016). These data have not been made publicly available, and the data in Pigeon et al. (2016) are embargoed until 2026. In addition, Festa-Bianchet and Pelletier (coauthors on Coltman et al. (2003) and Pigeon et al. (2016)) are signatories on Mills et al. (2015), which argues against making long-term individual-based data open access. Given it seems unlikely that these valuable data will not be made publicly available any time soon, we implore Coltman and colleagues to use their data to construct and analyze the class of model we use here. Until this is done, we recommend that the conclusions of Coltman et al. (2003) and Pigeon et al. (2016) are not used to inform wildlife management policies given their conclusions are not theoretically plausible.

Quantitative genetics theory is powerful, elegant, and based on irrefutable logic (Falconer, 1975; Lande and Arnold, 1983). The statistical methods used to estimate evolutionary change are also extremely powerful when assumptions that underpin the analyses are met (Lynch and Walsh, 1998). We recommend that when evolution is inferred from these statistical analyses, quantitative genetic theory based on the assumptions that underpin the analyses is used to check that reported patterns are plausible. For example, could a correlated character that results in the same selection differential that is observed on the trait generate the observed patterns? This is particularly important when statistically identified rates of evolution are very rapid, or occur in the opposite direction to that predicted. If patterns from these statistical analyses are not theoretically possible, some key assumption underpinning the statistical analysis has been violated, and conclusions from the statistical analyses are unreliable. Simple quantitative genetic models rarely provide predictions that match with observation in the wild (Merilä et al., 2001). When this happens, and predictions and observation cannot be reconciled, the use of phenotype-only models (Ellner et al., 2016), or models with more complex genotype-phenotype maps (Yang, 2004), can provide useful insight into causes of phenotypic change, particularly when these models capture observed dynamics accurately, as they frequently do (Coulson et al., 2010; Merow et al., 2014).

### Management Implications

Our work suggests that highly selective trophy hunting will result in evolutionary change, but that it will not be particularly rapid. Evolutionary change would be more rapid if both sexes were selectively targeted as is unfortunately the case for African elephant (*Loxodonta africana*) populations in some countries (Selier et al., 2014). When harvesting is less selective, or coupled with habitat change, the evolutionary consequences of selective harvesting may be harder to detect (Crosmary et al., 2013; Garel et al., 2007; Monteith et al., 2013; Rivrud et al., 2013). Our work does not tackle the ethics or ecological consequences of trophy hunting, nor do we account for potential economic benefits of hunting for local communities, whether these be in Canada (Hurley et al., 2015) or in the developing world (Lindsey et al., 2007). These issues should be given considerably more weight when designing population management and conservation strategies compared to the likelihood of rapid evolution.

## ACKNOWLEDGMENTS

We thank 2 anonymous reviewers for useful comments on an earlier version of this manuscript. TC also acknowledges support from the Natural Environment Research Council via a standard grant.

Associate Editor: Mark Boyce.

## Summary to the electronic Table of Contents

We show that the widely cited results claiming rapid evolution of trophy size in bighorn sheep in response to selective trophy hunting are theoretically impossible given standard assumptions of quantitative genetics.

